# Modelling cell type-specific lncRNA regulatory network in autism with Cycle

**DOI:** 10.1101/2024.05.31.594791

**Authors:** Chenchen Xiong, Mingfang Zhang, Haolin Yang, Xuemei Wei, Chunwen Zhao, Junpeng Zhang

**Affiliations:** School of Engineering, Dali University, 671000 Dali, Yunnan, China; Beijing CapitalBio Pharma Technology Co.,Ltd., 102200 Beijing, China; Beijing Computing Center, 100191 Beijing, China

## Abstract

Autism spectrum disorder (ASD) is a class of complex neurodevelopment disorders with high genetic heterogeneity. Long non-coding RNAs (lncRNAs) are vital regulators that perform specific functions within diverse cell types and play pivotal roles in neurological diseases including ASD. Therefore, studying the specific regulation of lncRNAs in various cell types is crucial for deciphering ASD molecular mechanisms. Existing computational methods utilize bulk transcriptomics data across all of cells or samples, which could reveal the commonalities of lncRNA regulation in the pathogenesis of ASD, but ignore the specificity of lncRNA regulation across various cell types. Here, we present *Cycle* (Cell type-specific lncRNA regulatory network) to construct the landscape of cell type-specific lncRNA regulation in ASD. We have found that each ASD cell type is unique in lncRNA regulation, and more than one-third and all of cell type-specific lncRNA regulatory networks are characterized as scale-free and small-world, respectively. Across 17 ASD cell types, we have discovered 19 rewired and 11 conserved modules, and eight rewired and three conserved hubs underlying within the discovered cell type-specific lncRNA regulatory networks. Moreover, the discovered rewired and conserved modules and hubs are significantly enriched in ASD-related terms. Furthermore, more similar ASD cell types tend to be connected with higher strength in the constructed cell similarity network. Finally, the comparison results demonstrate that *Cycle* is a potential method for uncovering cell type-specific lncRNA regulation.

## Introduction

Autism spectrum disorder (ASD) refers to a collection of neurodevelopmental disorders exhibiting profound genetic diversity and complexity [1,2]. Since childhood, ASD individuals have a wide range of difficulties and deficiencies in social interaction, language communication [3]. Despite striking progress in studying ASD has demonstrated that ASD possesses strong genetic heterogeneity and numerous molecules participate in regulating a series of complex biological processes, including neuronal activity [4] and immune response [2], an understanding of the pathobiology of ASD is still largely unclear. Unlocking the underlying pathogenesis of ASD at the molecular regulatory level holds profound implications in early detection and personalized treatment.

Long non-coding RNAs (lncRNAs) comprise a category of non-coding RNAs that are typically longer than 200 nucleotides, which act as regulators to make significant contributions to neurological diseases, e.g. ASD [3,5]. In the field of neurobiology, previous studies [3,6] have revealed that numerous lncRNAs exert biological functions specific to cell types, including neuronal differentiation, synaptic development, and plasticity [7]. In addition, lncRNA regulation also exhibits to be tissue-specific [8], and cell developmental-stage specific [9]. Due to the heterogeneity and complexity in the development of ASD, studying cell type-specific or dynamic lncRNA regulation could provide a new perspective for discovering potential therapeutic strategies for ASD.

At present, devising computational methods constitutes a highly promising way to decipher the function of lncRNAs in modulating ASD-related biological processes. By using bulk transcriptomics data, computational methods for identifying lncRNA regulation can be grouped into three primary categories: sequence-based methods that rely on nucleic acid sequence characteristics, expression-based methods focusing on variations in lncRNA expression levels, and integration-based methods that combine multiple sources of data. Sequence-based methods calculate the binding energy of RNA base pairs to infer lncRNA-target binding pairs. A prime example is LncTar [10], which utilizes the nearest neighbour thermodynamic model to compute the binding free energy of lncRNA-RNA pairs. Expression-based methods have been firmly established and encompass a diverse range of statistical [11,12], deep learning [13,14], or causal inference [15] approaches. These methods utilize gene expression profiles to derive and establish lncRNA-target correlation or causality pairs. Alternatively, integration-based methods [16,17] combine a variety types of data (e.g., sequence information and expression profiles), thereby enhancing the precision and reliability of lncRNA target prediction. The major limitation of the above methods using bulk transcriptomics data is that they ignore the heterogeneity of lncRNA regulation across various samples (cell lines or tissues). As single-cell and single-nucleus RNA sequencing technology continues to evolve, inferring lncRNA regulation with single-cell or cell type resolution opens a way to specifically explore lncRNA regulation applicable to unique cells or cell types in ASD. Regarding cell-specific gene regulation, CSN (Cell-Specific Network) method [18] pioneers the construction of cell-specific networks using single-cell transcriptome data. Subsequently, as an improvement of CSN, c-CSN [19], loc-CSN [20], and p-CSN [21] are also presented to infer conditional, local, and partial cell-specific networks, respectively. Specifically, for exploring cell-specific miRNA regulation, CSmiR [22] has also been developed to investigate single-cell level modulation of miRNA expression. In terms of regulation specific to individual cell types, scHumanNet [23] aims to generate specialized gene regulatory networks (GRNs) for individual cell types by leveraging the information contained in the HumanNet reference interactome and single-cell expression data. The recently developed scMTNI [24] method integrates single-cell multi-omics datasets to build GRNs specific to cell types across cell lineages. However, these cell-specific or cell type-specific regulation approaches prioritize primarily on transcription factor or miRNA regulation, rather than lncRNA regulation. To infer lncRNA regulation specific to biological conditions, CDSlncR [9] could infer lncRNA regulatory networks corresponding to distinct developmental states of the brain neocortex. Given that the pathogenesis of ASD involves a series of cell types and biological processes regulated by lncRNAs, thus it is crucial to study cell type-specific lncRNA regulation in ASD.

To explore the dynamic lncRNA regulation across various ASD cell types, we put forward a distinctively original method, *Cycle* (Cell type-specific lncRNA regulatory network), to model cell type-specific lncRNA regulatory networks in ASD. *Cycle* has two main contributions as follows. Firstly, contrary to considering all types of interactions encompassing those between mRNAs, between lncRNAs, and also between lncRNAs and mRNAs, *Cycle* primarily concentrates on identifying lncRNA-mRNA interactions. Secondly, taking the diversity and specificity of cells and cell types into consideration, *Cycle* identifies lncRNA regulatory networks specific to each cell type.

We have applied *Cycle* into single-nucleus RNA-sequencing (snRNA-seq) data of ASD brain tissues [25] for modelling the landscape of cell type-specific lncRNA regulation in ASD. Our research has found that each ASD cell type is unique in lncRNA regulation. Notably, over one-third of the cell type-specific lncRNA regulatory networks are scale-free, and all of them exhibit to be small-world. Among 17 ASD cell types, we have inferred 19 rewired modules and 11 conserved modules, along with eight rewired hubs and three conserved hubs leveraging these cell type-specific lncRNA regulatory networks. Importantly, these discovered rewired and conserved modules, and conserved hubs are significantly associated with ASD-related terms. Additionally, ASD cell types that are more similar tend to be strongly connected in the constructed cell similarity network. Finally, our comparison results suggest that *Cycle* represents a promising approach in elucidating cell type-specific lncRNA regulation.

## Results

### he landscape of cell type-specific lncRNA regulation in ASD

Following the workflow of *Cycle*, we model the landscape of lncRNA regulation across 17 ASD cell types. The number of lncRNA-mRNA interactions and hub lncRNAs tends to be various across 17 ASD cell types (**Figure 1a**). In the case of lncRNA-mRNA interactions, L4 and microglia cells obtain the largest and least number of interactions, respectively. In the case of hub lncRNAs, L2/3 and ASTFB have the largest and least number of hubs, respectively. Network topological analysis displays that 6 out of 17 (∼35.29%) cell type-specific lncRNA-mRNA regulatory networks adhere to a power law distribution, and all of cell type-specific lncRNA-mRNA networks display higher densities compared to their corresponding random networks (**Figure 1b** and **Supplementary Table 1**). These results indicate that over one-third of these cell type-specific lncRNA regulatory networks tend to be scale-free, and all of these cell type-specific lncRNA regulatory networks exhibit to be small-world.

**Figure 1:**
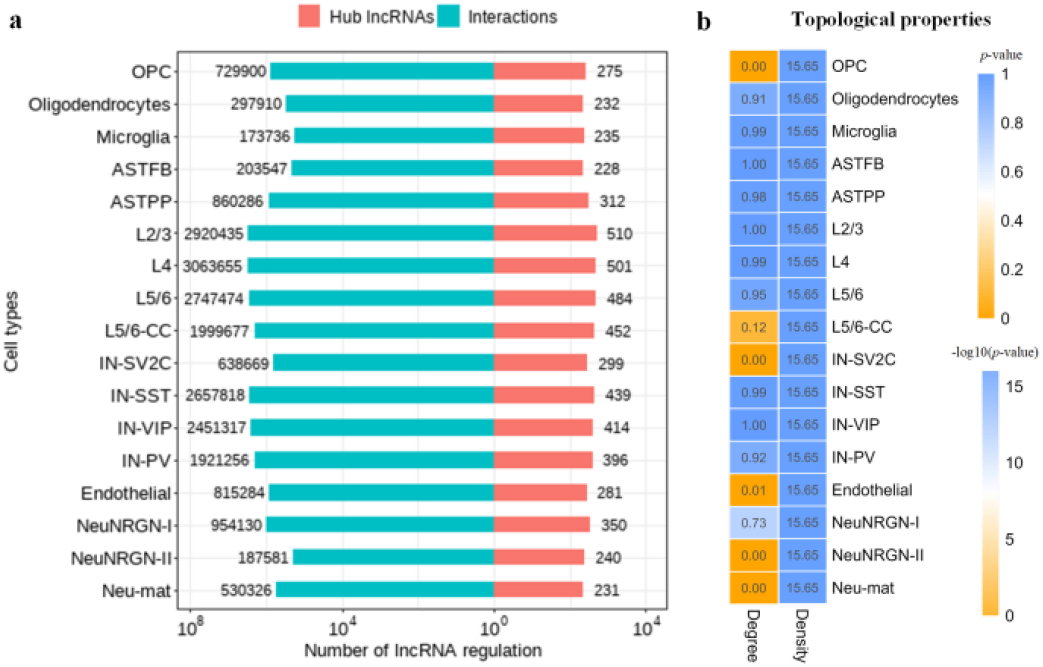
The lncRNA regulation landscape across 17 cell types. (a) The number of lncRNA-mRNA interactions and hub lncRNAs within each cell type. (b) The topological properties of lncRNA-mRNA regulatory networks for each cell type.

### Each ASD cell type is unique in lncRNA regulation

In this section, we investigate the uniqueness, conservation and dynamics of lncRNA regulation among various ASD cell types. We have found that the lncRNA regulatory networks and hub lncRNAs between any couples of 17 ASD cell types are various, indicating the uniqueness of each cell type (**Figure 2a** and **2b**). For the lncRNA regulatory networks, nearly half pairs between 17 cell types (∼46.32%) have a difference value with more than 0.500. Specifically, the highest difference value (0.951) between 17 cell types is between microglia and NeuNRGN-II (**Figure 2a**). For hub lncRNAs, more than one-third pairs between 17 cell types (∼36.76%) have a difference value with more than 0.500, and the highest difference value (0.844) between 17 cell types also exists between L5/6-CC and Neu-mat (**Figure 3b**). These results have suggested that each ASD cell type is unique in lncRNA regulatory networks and hub lncRNAs.

**Figure 2:**
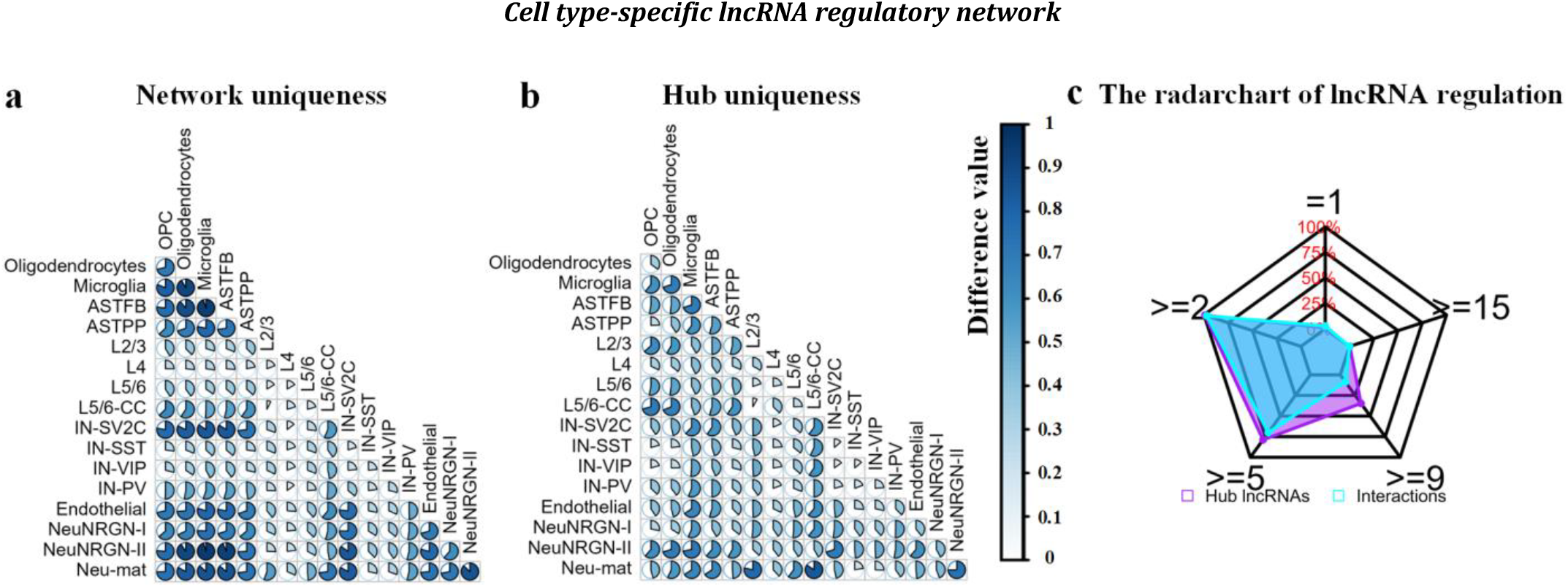
Uniqueness of lncRNA regulation across ASD cell types. (a-b) Uniqueness of lncRNA-mRNA regulatory networks and hub lncRNAs in each ASD cell type. (c) The radar chart of lncRNA-mRNA interactions and hub lncRNAs.

**Figure 3:**
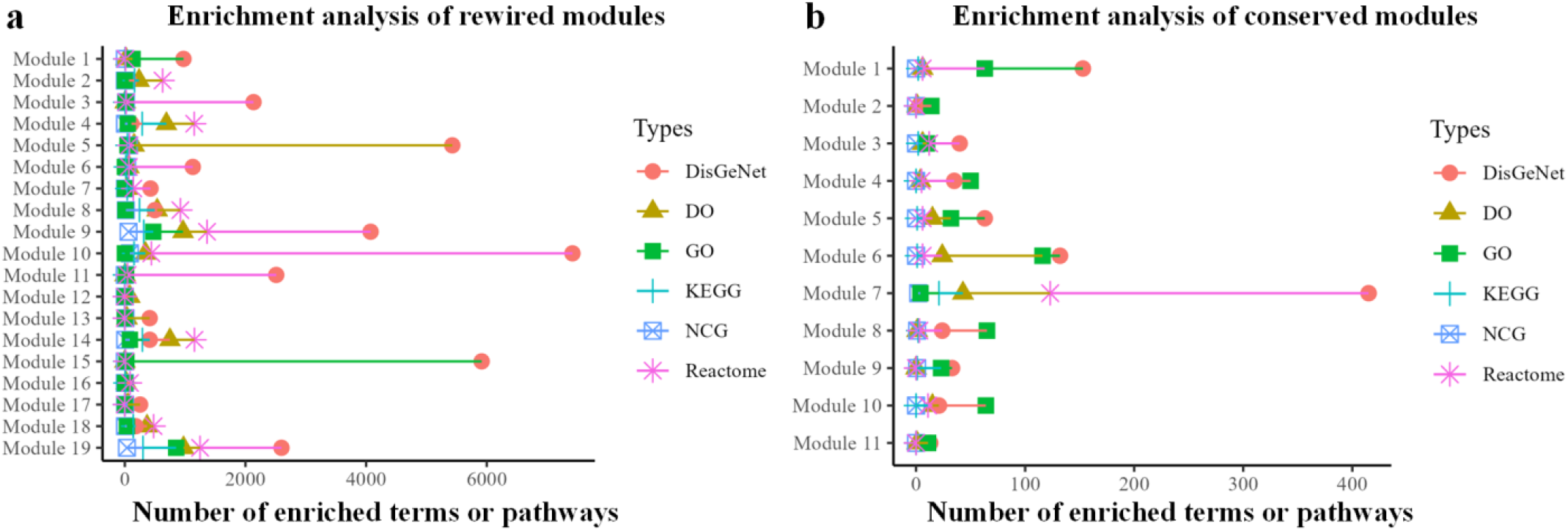
Enrichment analysis of rewired and conserved modules. (a) The number of enriched terms for rewired modules. (b) The number of enriched terms for conserved modules.

With regard to conservative and dynamic analysis, 115,108 lncRNA-mRNA interactions and eight hub lncRNAs (*CPVL-AS2, LINC00343, LINC01202, LINC01619, LINC01811, LINC02301, LINC03013, SYNPO2L-AS1*) only exist in one cell type, and 63 lncRNA-mRNA interactions and three hub lncRNAs (*ANKRD17-DT, LINC01572, MIRLET7BHG*) exist in at least 90% cell types (**Figure 2c**). In total, we have obtained 115,108 rewired interactions, 63 conserved interactions, eight rewired hubs, and three conserved hubs across 17 ASD cell types. Overall, the number of rewired interactions or hubs is larger than that of conserved interactions or hubs, indicating that lncRNA regulation tends to be dynamic across ASD cell types.

### Rewired and conserved lncRNA regulatory modules and hub lncRNAs are closely associated with ASD

Based on the rewired and conserved lncRNA-mRNA regulatory networks, we have further discovered 19 rewired and 11 conserved modules. To reveal the fundamental biological significance of lncRNA regulation associated with ASD, we conduct functional and disease enrichment analysis of the rewired and conserved lncRNA regulatory modules and hub lncRNAs. We have found that all of the rewired and conserved lncRNA regulatory modules exhibit significant enrichment in one or more terms belonging to GO, KEGG, Reactome, DO, NCG or DisGeNET databases (**Figure 3**). Among 19 rewired lncRNA regulatory modules, Module 10 has the largest number of terms or pathways enriched. For 11 conserved lncRNA regulatory modules, Module 7 has the highest number of enriched terms or pathways enriched. For the rewired and conserved lncRNA regulatory modules, several significant functional enriched terms, e.g., the GO term ″regulation of neuron projection development (GO:0010975)″, KEGG pathway “Pathways of neurodegeneration - multiple diseases (hsa05022)”, Reactome pathway ″Neuronal System (R-HSA-112316)″ show a close relationship with the progression and emergence of ASD traits. In addition, DisGeNET terms (Autistic behavior (C0856975) and Abnormality of brain morphology (C4021085), DO terms (autism spectrum disorder (DOID:0060041)), and NCG term (pan-cancer_paediatric) are also highly related to ASD (refer to **Supplementary Table 2** for more detailed information).

From the rewired and conserved lncRNA-mRNA regulatory networks, we have inference eight rewired hubs and three conserved hubs. The rewired hubs are not found to be significantly enriched in any particular biological pathway, and the conserved hub lncRNAs are significantly enriched in 184 KEGG, 3074 GO, and 364 Reactome terms or diseases, indicating their participate in ASD-related biological processes. The rewired hubs did not exhibit enrichment on the pathways from the functional analysis database. Notably, several of these terms, such as Reactome pathway ″PI3K/AKT Signaling in Cancer (R-HSA-2219528)”, KEGG pathway ″PI3K-Akt signaling pathway (hsa04151)”, and GO term ″negative regulation of neurogenesis (GO:0050768)”, are strongly linked to ASD (see details in **Supplementary Table 2**).

Altogether, the rewired and conserved lncRNA regulatory modules, and conserved hub lncRNAs are strongly linked with the pathophysiological progression of ASD.

### Cell similarity network

In this section, we further construct a cell similarity network by using lncRNA-mRNA interactions and hub lncRNAs in each ASD cell type. If the similarity value of a cell-cell pair is larger than the median value of similarity, the cell-cell pair is considered to be a link in the cell similarity network. As a result, we have found that L4 is similar with the largest number of other ASD cell types, while ASTFB is similar with the least number of other ASD cell types (**Figure 4**).

**Figure 4:**
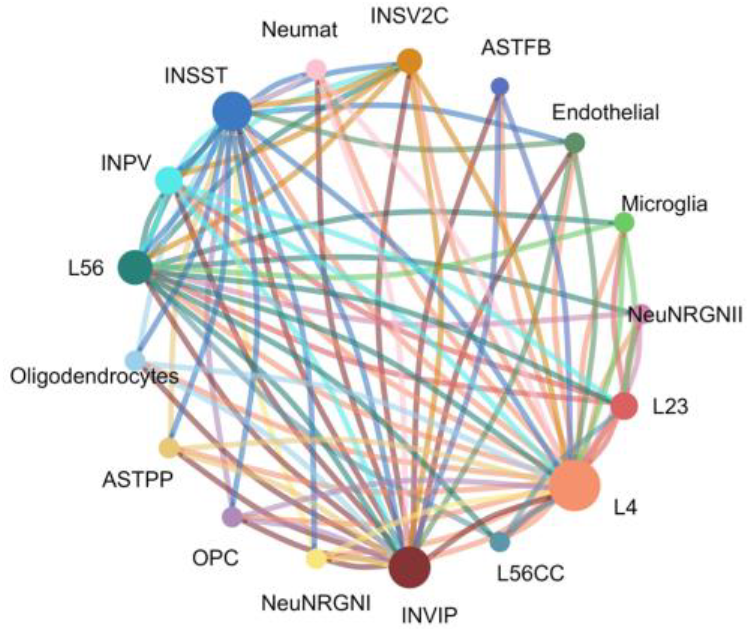
Cell similarity network. Cell similarity network between 17 ASD cell types by using lncRNA-mRNA interactions and hub lncRNAs. A larger circle denotes that the cell type is similar with a larger number of other cell types.

### In comparison with the other method

CDSlncR [9] is the first method to carry out research on cell type-specific lncRNA regulation. In this section, a performance comparison was conducted between *Cycle* and CDSlncR in terms of modelling cell type-specific lncRNA regulation. To ensure fairness, the significance threshold for *p*-value of CDSlncR and *Cycle* is set equally. We conduct a comparison of the number of validated lncRNA-mRNA interactions predicted by *Cycle* and CDSlncR [9]. For each ASD cell type, *Cycle* outperforms CDSlncR by yielding a higher number of validated lncRNA-mRNA interactions (**Figure 5**). The comparison yields insights indicating that Cycle is better than CDSlncR in modelling cell type-specific lncRNA regulation.

**Figure 5:**
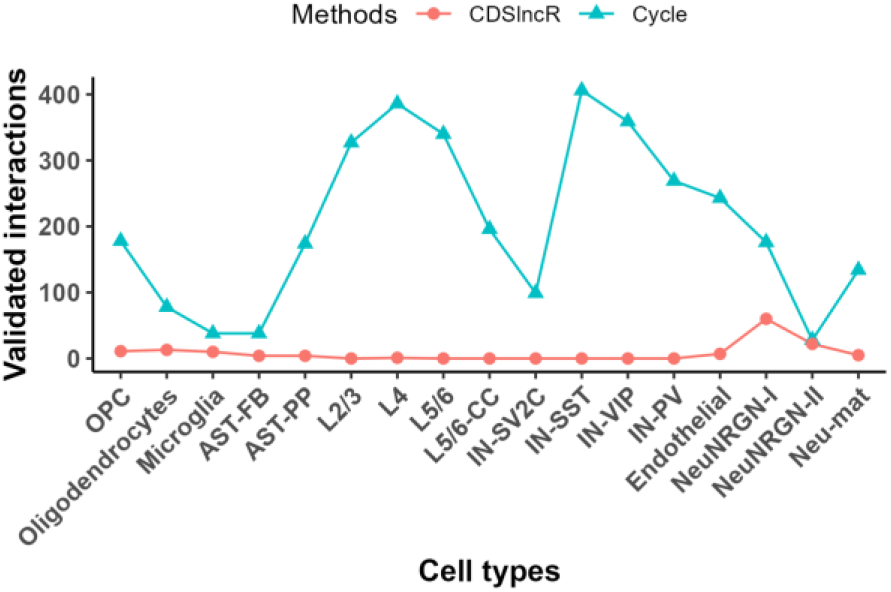
Comparison results. Comparison between *Cycle* and CDSlncR in the number of validated lncRNA-mRNA interactions.

## Materials and Methods

### The flowchart of Cycle

*Cycle* includes three main components (**Figure 6**). Firstly, *Cycle* conducts data preprocessing including gene annotation, feature selection, and data splitting to acquire the expression data pertaining to the highly expressed lncRNAs and mRNAs in 17 ASD cell types. For each ASD cell type, *Cycle* further identifies lncRNA regulatory networks specific to it. In total, 17 cell type-specific lncRNA regulatory networks are modelled. Derived from the constructed cell type-specific lncRNA regulatory networks, *Cycle* further deduces the rewired and conserved modules and hubs. Finally, *Cycle* can perform four types of downstream analyses, including network topological analysis, uniqueness analysis, cell similarity network construction, and enrichment analysis. The details of each component will be described in the following.

**Figure 6:**
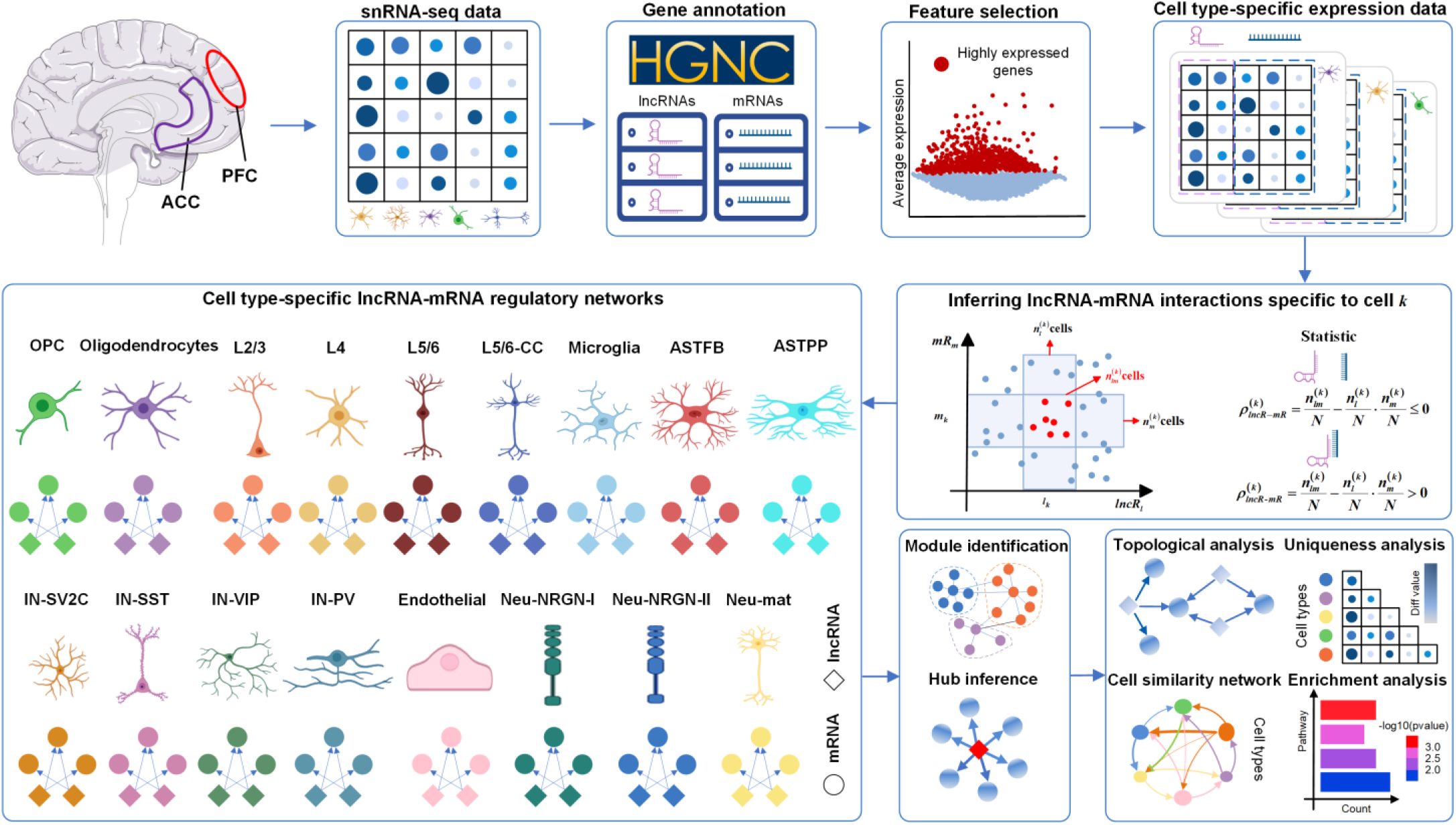
Workflow of *Cycle*. Firstly, *Cycle* extracts the matched lncRNA and mRNA expression data by using gene annotation information from HGNC (HUGO Gene Nomenclature Committee), and further retains the highly expressed lncRNAs and mRNAs for each cell type. In total, we have obtained 17 cell type-specific expression data of highly expressed lncRNAs and mRNAs. Secondly, *Cycle* models cell type-specific lncRNA regulatory networks for 17 ASD cell types. Furthermore, *Cycle* identifies the rewired and conserved modules, and infers hubs utilizing the established lncRNA regulatory networks specifically modelled for individual cell types. Finally, *Cycle* conducts four types of downstream analyses, including modules identification, hub inference, network topological analysis, uniqueness analysis, cell similarity network construction, and enrichment analysis. Created with BioRender.com.

### Single-nucleus RNA-sequencing data in ASD

We obtain ASD snRNA-seq data from the Sequence Read Archive (SRA) with accession number PRJNA434002 [25]. As a preprocessing step, we utilized gene annotation information from HGNC (HUGO Gene Nomenclature Committee, https://www.genenames.org/) and selected genes whose expression levels were higher than the average expression level across all cells. In total, we have retained 813 lncRNAs and 5,133 mRNAs highly expressed in 52,003 ASD cells. The 52,003 ASD cells are categorized into 17 cell types, including oligodendrocyte precursor cells (OPC), oligodendrocytes, microglia cells, fibrous astrocytes (ASTFB), protoplasmic astrocytes (ASTPP), layer 2/3 excitatory neurons (L2/3), layer four excitatory neurons (L4), layer 5/6 corticofugal projection neurons (L5/6), layer 5/6 cortico-cortical projection neurons (L5/6-CC), SV2C interneurons (IN-SV2C), somatostatin interneurons (IN-SST), VIP interneurons (IN-VIP), parvalbumin interneurons (IN-PV), endothelial cells, NRGN-expressing neurons (NeuNRGN-I), NRGN-expressing neurons (NeuNRGN-II), and maturing neurons (Neu-mat). Detailed information on 17 ASD cell types can be found in **Supplementary Table 1**.

### Identification of cell type-specific lncRNA regulatory networks

For each cell type, modelling cell type-specific networks is grounded upon the identification and characterization of cell-specific regulatory networks. Hence, the initial undertaking for the *Cycle* method is to precisely determine cell-specific lncRNA regulatory networks. Here, *Cycle* adapts CSN [18] with local strategy [20] to quantitatively estimate the correlation strength of lncRNA-mRNA relationship pairs in each cell. Within each cell, significantly cell-specific lncRNA-mRNA interactions are subsequently consolidated to model a cell-specific lncRNA-mRNA regulatory network.

In cell *k, l*_*k*_ and *m*_*k*_ are the expression values of lncRNA *lncR*_*l*_ and mRNA *mR*_*m*_, respectively, 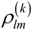 is calculated as the interaction strength between *lncR*_*l*_ and *mR*_*m*_ in the following:

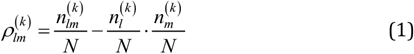

where*N* is the number of cells for ASD snRNA-seq data, 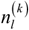 and 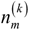 are the neighbourhood number of *l*_*k*_ and *m*_*k*_ in the bins of cell *k* for *lncR*_*l*_ and *mR*_*m*_, respectively, and 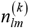 is the neighbourhood number of (*l*_*k*_, *m*_*k*_) in the interaction bin of cell *k*.

Owing to the specificity and heterogeneity of cells, self-adaptive window size 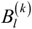 and 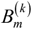 of bins in cell *k* are iteratively generated based on local standard deviations as follows:

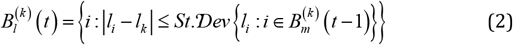

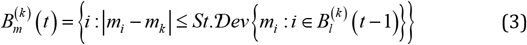

where initial *B*_*l*_ (0) and *B*_*m*_ (0) are a quantile range, and *t* starts from 1, 2,…, until convergence is achieved. If convergence is not achieved in the iterations, 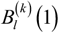 and 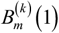 will be adopted as window sizes in practice for cell *k*.

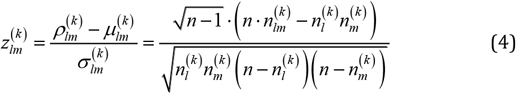

where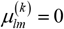 represents the mean value of 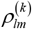, and 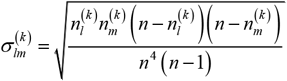 denotes the standard deviation of 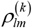. Every 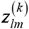 value carries a corresponding p-value, and a significantly smaller p-value (e.g., p < 0.01) evidences higher credibility for genuine lncRNA-mRNA interactions from a statistical perspective.

For each cell, we only focus on the lncRNA-mRNA interactions with statistical significance (e.g., p-value less than 0.01). If a significant lncRNA-mRNA interaction exists in more than 90% of total cells of a cell type, the lncRNA-mRNA interaction is regarded as one of a collection of lncRNA-mRNA interactions specifically for the cell type. By integrating all of the distinctive lncRNA-mRNA interactions peculiar to individual cell types, *Cycle* constructs 17 cell type-specific lncRNA-mRNA regulatory networks.

### Network topological analysis

Topological analysis contributes to exploring the characteristics and organization of biological networks including lncRNA regulatory networks. Degree and density are two widely used metrics to characterize a biological network. If the node degree distribution of a cell type-specific lncRNA regulatory network adheres to a power-law distribution with a *p*-value of a Kolmogorov-Smirnov test [26] larger than 0.05, the network tends to be a scale-free network. If the characteristic density of a cell type-specific lncRNA regulatory network is higher than that of its corresponding random networks at a significance level (e.g., 0.05), the network is regarded as a small-world network. Here, for each cell type-specific lncRNA regulatory network, we generate 1,000 random networks by randomizing the lncRNA-mRNA interactions. We utilize the Student’s t-test for statistically quantifying the differences between the built cell type-specific lncRNA regulatory networks and their corresponding random networks. In this work, the *igraph* R package [27] is applied to analyze the topological attributes of the constructed cell type-specific lncRNA regulatory networks.

### Hub lncRNA inference

Hub lncRNAs with high connectivity play key pivot roles in a cell type-specific lncRNA regulatory network. Rather than inferring hubs as those with a node degree exceeding a giving value, we assume that the node degree of lncRNAs follows the Poisson distribution [28–30]. For each lncRNA, we calculate the *p*-value of it accordingly presented:

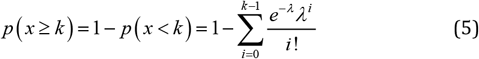

where*λ* = *np*, 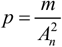, *n* is the number of lncRNAs, *m* is the number of lncRNA-mRNA pairs in a lncRNA–mRNA regulatory network, and 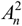 is the number of all possible lncRNA-mRNA interactions. In this work, a lncRNA with *p*-value less than 0.05 is viewed as a hub lncRNA.

### Conservative and dynamic analysis

In the cell type-specific lncRNA regulatory networks, the conserved and rewired interactions and hubs reveal the commonality and heterogeneity of different ASD cell types, providing new insights into conservative and dynamic lncRNA regulation across ASD cell types. Previous studies [15,31] have shown that lncRNA regulation is ‘on’ in some biological conditions but is ‘off’ in other biological conditions. Here, lncRNA-mRNA interactions or hub lncRNAs existing in at least 90% ASD cell types are considered as the conserved lncRNA regulatory network or hub lncRNAs, and lncRNA-mRNA interactions or hub lncRNAs existing in only one ASD cell type are viewed as the rewired lncRNA regulatory network or hub lncRNAs. To further identify highly connected functional modules within the conserved and rewired lncRNA regulatory networks, we have applied the Markov Cluster (MCL) algorithm to discover the conserved and rewired lncRNA regulatory modules. In every module, the combined total of lncRNAs and mRNAs should amount to at least three.

### Uniqueness of cell type-specific lncRNA regulation

In-depth exploration of the uniqueness of lncRNA regulation between ASD cell types involves focusing on the difference of lncRNA-mRNA interactions or hub lncRNAs between any pairs of cell types. Regarding cell type-specific lncRNA regulatory networks, we use the Simpson model [32] to estimate the similarity *sim*_*ij*_ between ASD cell types *i* and *j* . The difference *dif*_*ij*_ between ASD cell types *i* and *j* is computed as described below.

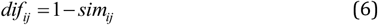

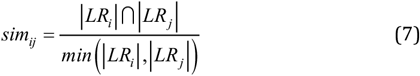

where*LR*_*i*_ and *LR*_*j*_ are lncRNA-mRNA interactions or hub lncRNAs existing in ASD cell types *i* and *j*, | *LR*_*i*_| ∩ | *LR*_*j*_| represents the intersection number of lncRNA-mRNA interactions or hub lncRNAs between *LR*_*i*_ and *LR*_*j*_, and *min* ( |*LR*_*i*_|, |*LR*_*j*_|) is the smaller number of lncRNA-mRNA interactions or hub lncRNAs between *LR*_*i*_ and *LR*_*j*_ . A larger value of *dif*_*ij*_ denotes a higher uniqueness between ASD cell types *I* and *j* .

### Enrichment analysis

To understand the conservative and dynamic biological processes in the conserved and rewired lncRNA-mRNA regulatory modules, we conduct enrichment analysis with *miRspongeR* [33] and *clusterProfiler* [34] R packages. The databases used for functional enrichment analysis include Gene Ontology (GO) [35], Kyoto Encyclopedia of Genes and Genomes (KEGG) [36], and Reactome Pathway database (Reactome) [37]. Additionally, three disease databases including Disease Ontology (DO) [38], DisGeNET [39], and Network of Cancer Genes (NCG) [40] are also considered for disease enrichment analysis. With regard to hub lncRNAs, we employ RNAenrich [41], a powerful comprehensive web server for ncRNA functional enrichment, to explore potential pathways, biological processes and diseases which they participate. In this work, the enriched KEGG, GO, Reactome, DO, DisGeNET or NCG term that exhibits a statistically significant enrichment with an adjusted *p*-value<0.05 (adjusted by the Benjamini-Hochberg approach) is viewed as notably enriched. Moreover, we collect experimentally validated lncRNA-target interactions from NPInter v5.0 [42], LncTarD v2.0 [43] and LncRNA2Target [44].

## Discussion

ASD is a set of complex neurodevelopmental disorders that manifest with varying symptoms among individuals. Exploring the regulatory mechanisms of lncRNA within and between different ASD cell types holds significance in elucidating the etiology and ontogeny of ASD. Our work develops a novel approach called *Cycle*, designed to model cell type-specific lncRNA regulatory networks in ASD. For each ASD cell type, we have shown that the lncRNA regulation tends to be unique. Moreover, the rewired and conserved lncRNA regulatory modules and hub lncRNAs are significantly enriched in several ASD-related terms or pathways. In addition, cell similarity network can help to know which cell types are similar with the least or largest number of other ASD cell types. In comparison with CDSlncR, *Cycle* performs better in inferring cell type-specific lncRNA regulation.

In future, *Cycle* can be further improved in the following three aspects. Firstly, *Cycle* mainly focuses on lncRNA regulation specific to ASD cell types. In future, it is necessary to study condition-specific lncRNA regulation, e.g., sex-specific or region-specific lncRNA regulation. Secondly, *Cycle* only infers the association/correlation rather than causal relationships between lncRNAs and mRNAs. In future, we will conduct cell type-specific lncRNA causal regulation research. Thirdly, competing endogenous RNA (ceRNA) hypothesis [45] suggests that lncRNAs have the potential to modulate gene expression by acting as ceRNAs, thus it is strongly needed to identify cell type-specific lncRNA-related ceRNA networks for comprehensively understanding lncRNA regulation.

## Conclusion

Overall, *Cycle* is useful for modelling the landscape of cell types-specific lncRNA regulation in ASD. *Cycle* gains insights into the lncRNA regulation underlying ASD across various cell types, and provide potential treatment strategies.

## Acknowledgements

This work has been supported by the National Natural Science Foundation of China (Grant Number: 61963001), the Yunnan Xingdian Talents Support Plan—Young Talents Program, and the Applied Basic Research Foundation of Science and Technology of Yunnan Province (Grant Number: 202101BA070001-221).

## Author contributions

Chenchen Xiong: Conceptualization, Data curation, Formal analysis, Methodology, Software, Visualization, Writing – original draft; Mingfang Zhang: Formal analysis, Software, Resources; Haolin Yang: Conceptualization, Resources, Writing – review & editing; Xuemei Wei: Conceptualization, Resources, Writing – review & editing; Chunwen Zhao: Conceptualization, Resources, Writing – review & editing; Junpeng Zhang: Conceptualization, Data curation, Formal analysis, Methodology, Validation, Funding acquisition, Writing – review & editing. All authors reviewed the manuscript.

